# Unraveling the Mysteries of Sleep: Exploring Phylogenomic Sleep Signals in the Recently Characterized Archaeal Phylum Lokiarchaeota near Loki’s Castle

**DOI:** 10.1101/2024.11.01.621620

**Authors:** Seithikurippu R Pandi-Perumal, Konda Mani Saravanan, Sayan Paul, David Warren Spence, Saravana Babu Chidambaram

## Abstract

Sleep is a universally conserved behavior with an elusive origin and an uncertain evolutionary purpose. Leveraging phylogenomics, we investigate the evolutionary foundations of sleep by identifying orthologs of Human sleep-related genes in the Lokiarchaeota of the Asgard superphylum. Our findings indicate a conserved suite of genes associated with energy metabolism and cellular repair, suggesting a primordial role of sleep in cellular maintenance. This data lends credence to the idea that sleep improves organismal fitness across evolutionary time by acting as a restorative process. Notably, our approach demonstrates that phylogenomics is more useful than standard phylogenetics for clarifying common evolutionary traits. By offering insight into the evolutionary history of sleep and putting forth a novel model framework for sleep research across taxa, these findings contribute to our growing understanding of the molecular foundation of sleep. This study lays the groundwork for further investigations into the importance of sleep in various organisms, which could have consequences for human health and a deeper comprehension related to the fundamental processes of life.

**Graphical Abstract:** 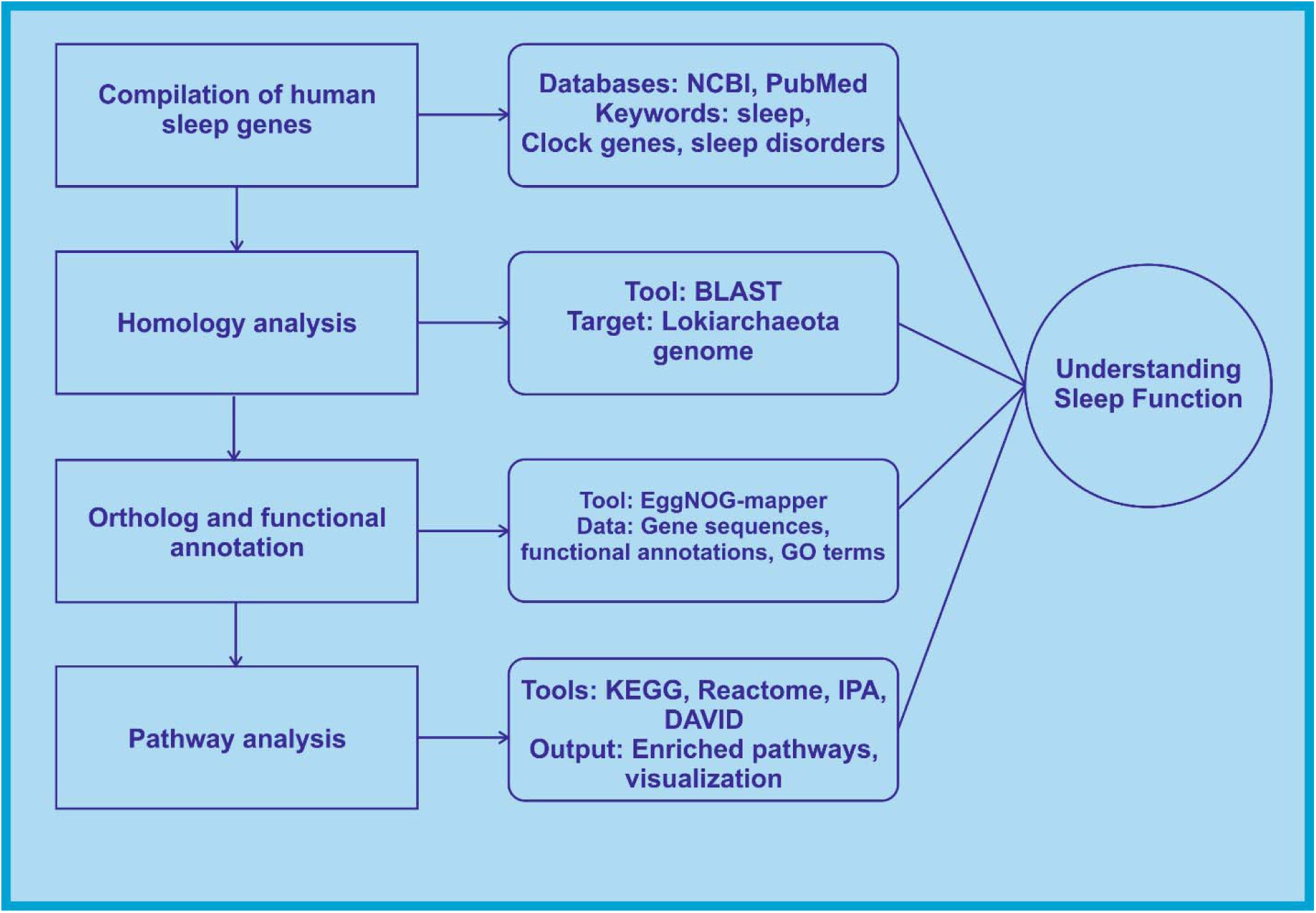

## Introduction

Since antiquity, the concept of sleep has been mentioned in literary or religious writings. Most ancient civilizations recognized and acknowledged the nature and value of healthy sleep in their lives, and sometimes even identified the gods who influenced or even caused sleepers’ dreams. For example, sleep and its connections to the supernatural world are focal topics in the Sumerian "Epic of Gilgamesh" (circa 2100 BCE) as well as in the Rigveda of the ancient Hindu and Vedic literature (circa 1500 and 1200 BCE or even older). Sleep and dreams were subsequently viewed from many perspectives for centuries (BaHammam et al, 2018; Kumar, 2015; Heydari et al, 2013; Barbera, 2008). The topic has been considered by novelists, playwrights, statesmen, and scientists, all of whom have contributed their particular interpretive wisdom. In the 19^th^ and 20th centuries, however, the general conclusions of these efforts were more systematically explored by the modern field of sleep medicine (lshimori K, 1909; Howell, 1913). The empirical method was applied, in many cases for the first time, to study the general principles underlying the sleep process in both humans and animals.

However, it was the discovery of the phenomenon of rapid eye movement (REM) sleep in 1953 which was a watershed event that elevated sleep studies into a major scientific endeavor, and marked the beginning of modern sleep research (Aserinsky & Kleitman, 1953). Over the decades, these studies have spawned numerous hypotheses and theories about sleep, and the list keeps expanding (Pandi-Perumal, 2010; Rechtschaffen, 2015; Moorcroft, 1995). Despite these advances, several information gaps still exist in terms of its organization, process, mechanism, regulation, and function, and thus our knowledge of sleep processes is still incomplete. Some of these theories and hypotheses, for instance, are speculative, have a restricted scope (partial explanation), don’t offer any concrete proof (negative findings), and don’t apply to all species. Consequently, most of these theories are either too narrowly focused or too divergent, or they are contradictory. Additionally, some of their associated ideas about sleep have been debated or refuted by contemporary scientists (Frank, 2013; Tononi & Cirelli, 2012; Rial et al, 2012, 2007; Rattenborg et al, 2008; FRANK & CRAIG HELLER, 2005; Vertes, 2004; Siegel, 2001; Frank & Heller, 2003; BLUMBERG et al, 2005; Siegel, 1995). Readers are referred to these reviews for a more in-depth consideration of these controversies and arguments.

"If sleep does not serve an absolutely vital function, then it is the biggest mistake the evolutionary process has ever made," according to Allan Rechtschaffen. This insightful assertion suggests that if sleep were a pointless behavior, evolutionary pressure may have prevented it from emerging, which is not the case. Nevertheless, despite the obvious relevance of sleep in sustaining the life process, there continues to be disagreement regarding its characteristics and components (Rial et al, 2007, 2012; Geissmann et al, 2024). Even if its exact purpose is unknown, sleep is something we all do for roughly one-third of our lives, an existential fact that suggests that sleep obviously has some critical, life-supporting function.

What happens if conventional methods or ways of thinking fail to answer longstanding questions in biology and neuroscience, such as why do we sleep? Inasmuch as “tried and true” techniques of investigation have failed to resolve these basic questions about sleep, perhaps some alternative approaches would merit consideration. It is hoped that a combination of the latest methods drawn from phylogenomics and bioinformatics methods may productively focus the hunt for the primordial function of sleep (Pandi-Perumal et al, 2024; Eisen & Fraser, 2003). In pursuit of this objective, we have endeavored to examine this core issue from a standpoint that has not been previously examined.

### Rationale for the chosen experimental approach

Our journey to develop this hybrid strategy of combining phylogenomics and bioinformatics begins with the study of unusual marine microorganisms that have only been found in remote underwater regions. These difficult-to-access locations host hydrothermal vents which in turn contain a rich but only recently characterized class of living organisms. One of these locations is known as Loki’s Castle. This site (sometimes referred to as Loki’s Castle Vent Field [LCVF]) was first discovered in 2008 and is approximately 2,300 meters below the surface of the water. It is located halfway between Greenland and Norway, near the intersection of the Mohns and Knipovich Mid-Atlantic Ridges (Pedersen et al, 2010). This region contains a field of so-called hydrothermal vents, i.e., outcroppings which appear to be rocky, black chimneys. These vents spew out hot fluid discharges (maximum temperature is about 320° C) which originate from even deeper regions of the earth’s mantle. The makeup of these discharges reflects the diverse geological and geochemical composition of their constantly shifting sources (Sahlström et al, 2023). The age of these formations continues to be speculative, but analysis of the hydrothermal plume fallout has suggested that LCVF vent activity began about 10,000 years ago. Currently, the area is intensely dynamic, showing extreme fluctuations in temperature, pH, and toxic compounds. This activity contains multiple targets for scientific inquiry. The complexity of the geochemical composition of the vents, while representing a major challenge for investigative analysis, is nevertheless overshadowed by the even greater zoological mystery of how the associated floral and faunal life, which is diverse, has managed to survive in this harsh environment (Eilertsen et al, 2024; Jaeschke et al, 2012). The use of the term “life” in LCVF primarily relates to microorganisms such as bacteria and archaea., which have taken advantage of the chemically rich content from hydrothermal vents as their source of energy rather than sunlight. Consequently, these organisms have developed a tolerance to extreme environmental conditions. Unlike those on the planet’s dry surface, these include, e.g., a lack of sunlight, extremely high temperatures, and high levels of toxicity. Additionally, these microorganisms have made dietary adaptations such as obtaining nutrients from chemosynthetic microbes. Taken together, these internal adaptations and environmental stresses have produced in these microorganisms several cellular and organismal specializations.

One class of these microorganisms is that of Archaea, and, already in the accelerated evolutionary conditions of the LCVF, it has developed into various lineages, each with its anatomical peculiarities and life-supportive strategies. While originally Archaea were similar to bacteria in size and shape, they currently differ from bacteria in several ways: a. their cell walls contain pseudo peptidoglycan and not peptidoglycan as in the case of bacteria; b. their cell membrane contains ether-linked lipids as opposed to lipids in bacteria. More recent research has confirmed that Archaea have more traits in common with eukaryotes than with bacteria. For example, some proteins, such as the DNA-dependent RNA polymerase, were exclusively discovered in Archaea and Eukarya, to the exclusion of bacteria, and others were more closely related to their eukaryotic homologs (van Wolferen et al, 2022; Hatano et al, 2022). The archaeal RNA polymerase closely resembles eukaryotic RNA polymerases. Eukarya and Archaea possess multi-subunit, DNA-dependent RNA polymerases with considerable structural and sequence similarities (Best & Olsen, 2001). This similarity differs from the more basic, single-subunit bacterial RNA polymerase, suggesting that Eukarya and Archaea possess a more recent common ancestor and that complex RNA polymerases evolved singularly. The transcription apparatus of Archaea resembles that of eukaryotes, encompassing analogous transcription factors and promoter recognition mechanisms (Rivas & Fox, 2024). It illustrates the evolutionary pressures linked to the regulation of intricate genomes. These commonalities indicate the evolutionary divergence of Archaea and Eukarya from simpler microorganisms (Londei, 2005).

### Loki genome characteristics

Spang et al. (2015)(Spang et al, 2015) conducted a metagenomic investigation of marine sediment obtained about fifteen kilometers away from Loki’s Castle, an active hydrothermal vent in the Arctic Mid-Ocean Ridges (AMOR; 73.763167 N, 8.464000 E) that are ∼4000 kilometers long, located at 3283 m below sea level (Figure 1; adapted from [Fredrik Sahlström et al., 2023])(Sahlström et al, 2023), which resulted in the discovery of a novel archaeal phylum, Lokiarchaeota (low-key-ar-kay-oh-tuh or Lokiarchaeia; henceforth referred to as Loki), which are the closest living relatives of all eukaryotes. Later, it was described as the earliest known member of the Asgard superphylum (MacLeod et al, 2019).

**Figure 1.**
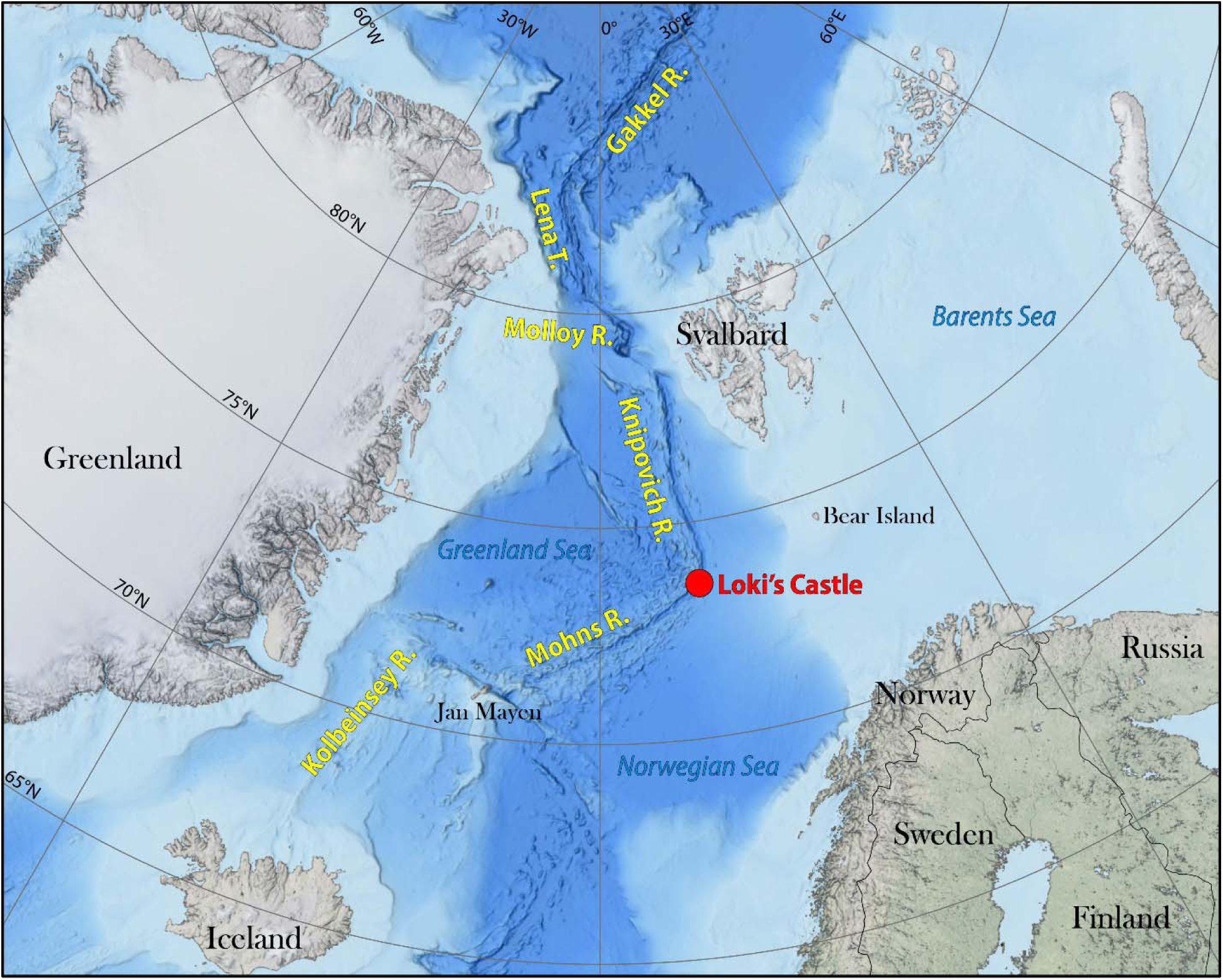
Bathymetric map showing the location of Loki’s Castle (3283 m below sea level with the following coordinates: 73.763167 N, 8.464000 E). The authors acknowledge Dr. Fredrik Sahlström for the supply of the image. The location from which the organism was collected is located 1.5 km north of Loki’s Castle. This location should still be within the red dot depicted in this image. Figure modified from Sahlström et al. (2023).

The Asgard phylum of archaea includes Lokiarchaeota, the study of whose genomes have contributed significantly to our understanding of the evolution of archaea from eukaryotes. In comparison, the genomes of Lokiarchaeota are notably larger when compared to other archaeal genomes and possess a diverse array of genes, including those related to the cytoskeleton, vesicular transport, and ubiquitin systems, which exhibit eukaryotic characteristics (MacLeod et al, 2019). These characteristics suggest that Lokiarchaeota could represent a crucial missing link between prokaryotic organisms and eukaryotic cells. Furthermore, these organisms exhibit significant metabolic versatility, with genes related to carbon fixation, sulfur compound metabolism, and hydrogen production. This indicates their potential to thrive in various environments despite their predominant association with marine sediments (Spang et al, 2015). The Scanning Electron Microscope (SEM) image of Ca. L. ossiferum reveals its intricate structure including morphology and its cellular architecture, characterized by coccoid-shaped cells with protrusions (Figure 2) (Rodrigues-Oliveira et al, 2023). This comprehensive observation indicates that this archaeal species located in hostile marine environment possesses morphological traits distinct from that of other bacterial species.

**Figure 2:**
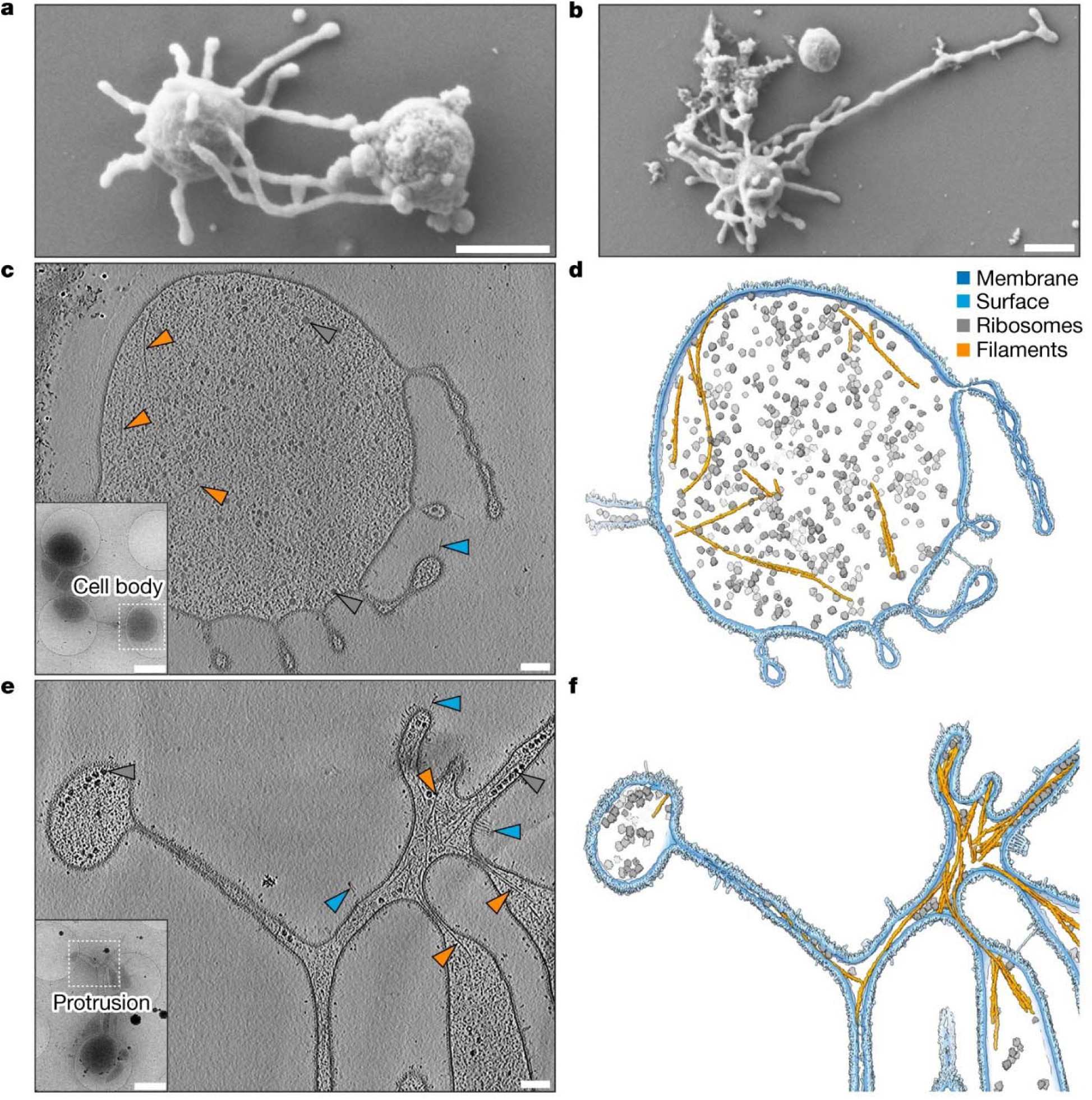
Complex and cellular anatomy of *Candidatus Lokiarchaeum ossiferum* cells. A scanning electron micrograph (SEM) of a *Ca. L. ossiferum* cell showing coccoid cells with long, complex cell protrusions. The image obtained at a high resolution defines detailed morphological and anatomical features that distinguish this type of archaeal species that inhabits marine and extremophiles’ environments. Such morphology found in this image helps in explaining the functional aspects and evolutionary lineage of *Ca. L. ossiferum* within the domain of Archaea. This image was taken with a scanning electron microscope and offers a close picture of the cell membrane, which is relevant to the analysis of microorganisms’ variation and evolution. (Adapted from, Rodrigues-Oliveira et al. Nature volume 613, pages 332–339 (2023). https://doi.org/10.1038/s41586-022-05550-y; licensed under <u>CC BY 4.0</u>).

The size of the genomes in Lokiarchaeota (4 to 5.5 megabases [Mb]) was unexpected, inasmuch as most archaeal genomes typically fall within the smaller range of 1 to 3 Mb (Zaremba-Niedzwiedzka et al, 2017). The larger size of the Lokiarchaeota genome is primarily attributed to the presence of genes associated with various eukaryotic-like processes, including actin, tubulin, small GTPases, and specific components of endosomal sorting complexes. The relevant genes are associated with more intricate cellular functions, such as cytoskeleton dynamics and vesicular motility, which are not widely observable in most archaea (Zaremba-Niedzwiedzka et al, 2017). The genome’s substantial size indicates the organism’s metabolic potential, encoding numerous genes related to the utilization of carbon, sulfur, and other compounds, along with potential environmental interactions. This genomic complication indicates that Lokiarchaeota may have served a significant evolutionary function as a transitional phase in the evolution of eukaryotic cells (Farag et al, 2020).

The Lokiarchaeum genome contains about 5381 proteins, of which 175 are signature proteins of eukaryotes. These proteins are involved in vital processes such as intracellular trafficking and transport, cytoskeletal activity, and cell membrane remodeling (Nasir et al, 2015). Some of these proteins were either absent or poorly expressed in the genomes of archaea and bacteria before. Taken together, this evidence further points to Lokiarchaeota’s potential status as a unique transitional link between other prokaryotic and eukaryotic cells.

## Methods

### Collection of Sleep Genes

In our ongoing phylogenomics and bioinformatics study, the process of identifying sleep-related genes initially involved conducting a keyword search on the NCBI Gene database (Sayers et al, 2024). The search focused on key terms such as sleep, circadian rhythm, clock genes, insomnia, sleep disorders, sleep regulation, and sleep genes. The data collected were exclusively applicable to Homo sapiens and were organized based on both the date and the type of investigation. Every outcome was condensed to infer findings on the stated genes while also taking note of any literature on sleep processes or disorders in the annotation. The research publications supplied the standard gene symbol, full name, and function in marginal for each gene. To confirm the involvement of these genes in sleep, an additional search was conducted in the PubMed database for each of the symbols and names to obtain relevant materials (Motschall & Falck-Ytter, 2005). This was also utilized to verify their participation in the regulation and problems of sleep, as shown in the GeneCards database (Barshir et al, 2021). To ensure the significance and validate the accuracy of the acquired gene list, supplementary sources such as Online Mendelian Inheritance in Man (OMIM) or the most recent sleep-related studies and reviews were consulted (Hamosh et al, 2021). Consequently, by systematically following the procedures outlined above, a comprehensive inventory of genes associated with sleep functions was successfully compiled for this and other ongoing studies. These discoveries served as the foundation for subsequent phylogenomic and bioinformatics investigations into the relationship between the orthologs identified in the species of interest to sleep genes.

### Blast analysis of sleep genes against the Lokiarchaeota genome

To conduct a Basic Local Alignment Search Tool (BLAST) analysis of sleep genes against the genomes of Lokiarchaeota (Spang et al, 2015), it was necessary to obtain the nucleotide or protein sequences of the sleep-associated genes from the National Center for Biotechnology Information (NCBI) in the Fast Approximation of Smith & Waterman Algorithm (FASTA) file format (Altschul et al, 1997). At the NCBI BLAST website, an appropriate BLAST-type query: the nucleotide for DNA sequences or protein for amino acid sequences was employed. The NCBI genome gateway is used to access the Lokiarchaeota genome. To enhance the alignments, the following settings were modified: the anticipated threshold for E-value, match/mismatch scores, and gap penalties. We performed a BLAST search to evaluate the hits based on the alignment scores, E-value, and percentage identity to identify the homologous sequences. Finally, to determine the biological importance of the similarities indicated earlier, it was necessary to compare the genes under examination with other sources of information. This comparison helped to clarify the function of the analyzed genes with the comparable genes found in Lokiarchaeota. Additionally, to ensure the preservation of the BLAST parameters, a concise and comprehensive description of each type of result, and visual representations of alignments was verified. Such an approach helped to observe the evolutionary conservation and functionality of different sleep genes across varied species.

### ORTHOLOGY AND FUNCTIONAL ANNOTATION (EGGNOG) OF SLEEP GENES

Prior to starting the annotation process of genes associated with sleep using the Evolutionary Genealogy of Genes: Non-supervised Orthologous Groups (EggNOG), the sleep genes’ protein or nucleotide sequences were first acquired in Fasta format from the NCBI. The EggNOG website was used to retrieve data from the EggNOG database, and the EggNOG-mapper was employed for annotation purposes. The sleep gene sequences were uploaded directly into the EggNOG-mapper interface as FASTA files (Huerta-Cepas et al, 2017). To attain optimal outcomes, the appropriate taxonomic level was selected, and additional factors, including the E-value, were adjusted. Sequence annotation proceeded similarly, with the significant reference set being the EggNOG orthologous groupings. The functional annotations were derived from conserved domains along with any other functionally related information for the implicated genes.

Pathways from the Kyoto Encyclopaedia of Genes and Genomes (Kanehisa & Goto, 2000), Clusters of Orthologous Genes (Makarova et al, 2007), and Gene Ontology keywords were used to further subclassify all sleep genes and provide functional descriptions for them. To establish clear connections between these genes and ongoing sleep processes and provide precise functional categorization, these annotations were meticulously compared to the most recent data available in the field of sleep science, ensuring the connections’ validity. Together with the comprehensive output from the tool EggNOG-mapper settings, a broad description of the genes with matching sequences co-annotated with EggNOG is provided.

### INTERPRO SCAN OF SLEEP GENES

Prior to commencing the EggNOG annotation of sleep genes, it was necessary to retrieve the nucleotide or protein sequences of the sleep genes from the NCBI database in FASTA file format (Blum et al, 2021). To acquire the database, the EggNOG website was navigated, and the EggNOG-mapper application was initiated. The sleep gene sequences were submitted to the EggNOG-mapper interface either by uploading the file or by manually copying and pasting the sequences. To minimally impact the outcomes, the most suitable taxonomic level was selected and other parameters, such as the E-value, were adjusted. Additionally, sequence annotation was performed using the EggNOG mapper after aligning it to EggNOG orthodox groups. This was followed by functional annotations based on conserved domains and other functional information. Subsequently, the consequences of functional descriptions obtained from Go words, kegg pathways, and COG for each sleep gene were analyzed. To validate these annotations using the latest research findings in the sleep field and establish a correlation between these genes and the ongoing sleep processes, it was necessary to give them appropriate functions. A comprehensive list of gene sequences that are co-annotated with EggNOG was provided. Additionally, detailed information about the settings utilized in the EggNOG-mapper tool, a description of the functional annotations, and graphical representations of pathways and biologically biased distributions of Gene Ontology (GO) terms were included. This technique provided comprehensive functional descriptions and biological function information regarding sleep-related genes.

### PATHWAY ANALYSIS OF SLEEP GENES

Before running the actual pathway analysis of sleep genes, it was necessary to have a list of sleep genes together with their gene codes in the form of gene symbols or Entrez numbers. Pathway analysis tools such as KEGG, Reactome, Metacore/Ingenuity Pathway Analysis (IPA), and DAVID were selected for further data processing (Kanehisa & Goto, 2000; Milacic et al, 2024). In doing so, one will be redirected to the software’s website. The gene list was entered in the appropriate format as directed by the tool. To control the analysis parameters, the organism of interest and the kind of analysis were set; this was either a pathway or a network. Pathway analysis was required to find out which of the presented pathways are affected and to what extent based on the genes. Looking at the enriched pathways next, highlighted genes and the significance levels of these genes were reviewed. To evaluate the role of the identified pathways in the biology of sleep, we validated the findings concerning the research work available in the literature. A specific report containing information about input genes was prepared, including parameters used in further analysis, the list of enriched pathways, and the significance of the obtained results containing visualizations in the form of pathway maps and network plots. This systematic approach offered the most efficient means for investigating the molecular aspects of sleep regulation and utilization.

## Results

### Orthologs of sleep genes in Lokiarchaeota

Table 1 presents a comparative BLAST study of sleep genes against the Lokiarchaeota genome. It was observed that many sleep genes showed substantial e-value and similarity scores when compared to Lokiarchaeota. The e-value (expected value) is crucial in BLAST to score the alignment outcomes. The term "e-value" refers to the likelihood of achieving such a score by chance; a low e-value indicates that the match is more likely to be biologically significant than an accidental event because similar sequences originate from related evolution. However, a high e-value suggests that the match may result from random similarities in the protein space, making it less valuable for additional biological research. Consequently, when examining homologous sequences or functional relatedness, researchers frequently consider the sequences with an e-value threshold (Chojnowski & Bochtler, 2007). The repertoire of BE- impacting proteins is extensive and varied, likely playing a crucial role in vital biological processes and the control of sleep. Interestingly, proteins such as the “molecular chaperone DnaK" and "ATP citrate synthase" show high similarity scores, 70. 27, and 73. 55, respectively, with the e-value zero, which suggests it is a solid match. These proteins are engaged in cellular stress and metabolisms that could be linked in one way or the other to sleep regulation (Table 1).

Examples of enzymes found include “4Fe-4S dicluster domain-containing protein", amidase, aldo/keto reductase, and diverse aminotransferases. These enzymes are involved in metabolic activities and redox reactions, which implies a possibility of links between metabolic management and sleep. The fact that there are many types of aminotransferases and reductases only highlights the relevance of metabolic and redox-regulating systems in the setting of sleep. The gene hits thus obtained like "ABC transporter ATP-binding protein" and the particular "protein kinase" are protein targets that are connected to signal transduction or transport processes in a variety of biological systems. These protein-coding genes are dominant in cells and have fundamental motifs that are consistently studied in molecular biology research and indicate the importance of certain proteins in basic cell processes. These functions are essential for the cellular signaling and regulation of other crucial processes that are important for the regulation of sleep.

“50S ribosomal protein L3”, elongation factor EF-2, and various transferases, including glutamine synthetase and NAD-dependent protein deacylase, assert the importance of protein synthesis and modification during sleep. There is a fundamental need for protein synthesis and modification to enable the cell to carry out its duties during rest, which is when it is sleeping. Additionally, our analysis showed multiple ‘hits’ to the ‘hypothetical proteins,’ which offer fascinating implications for the putative new function of sleep. The best way to define the "putative new function" of sleep is as a relatively recent concept that suggests sleep may have additional molecular functions not previously recognized; these may be tied to hypothetical proteins discovered in specific recent analyses. This new hypothesis suggests that sleep, particularly deep sleep, may be essential at a more basic molecular level for cellular repair or regeneration of DNA or even strengthening of the immune system. This new hypothesis adds to the conventional functions of sleep, such as restoration, energy conservation, and even optimization of cognitive processes. Although it could assume some significant sleep-related goals, this idea is mainly theoretical and based on molecular biology and phylogenomics developments. These chemicals remain to be described; consequently, they present promise for further study into the part they play in sleep regulation (Figure 3).

**Figure 3.**
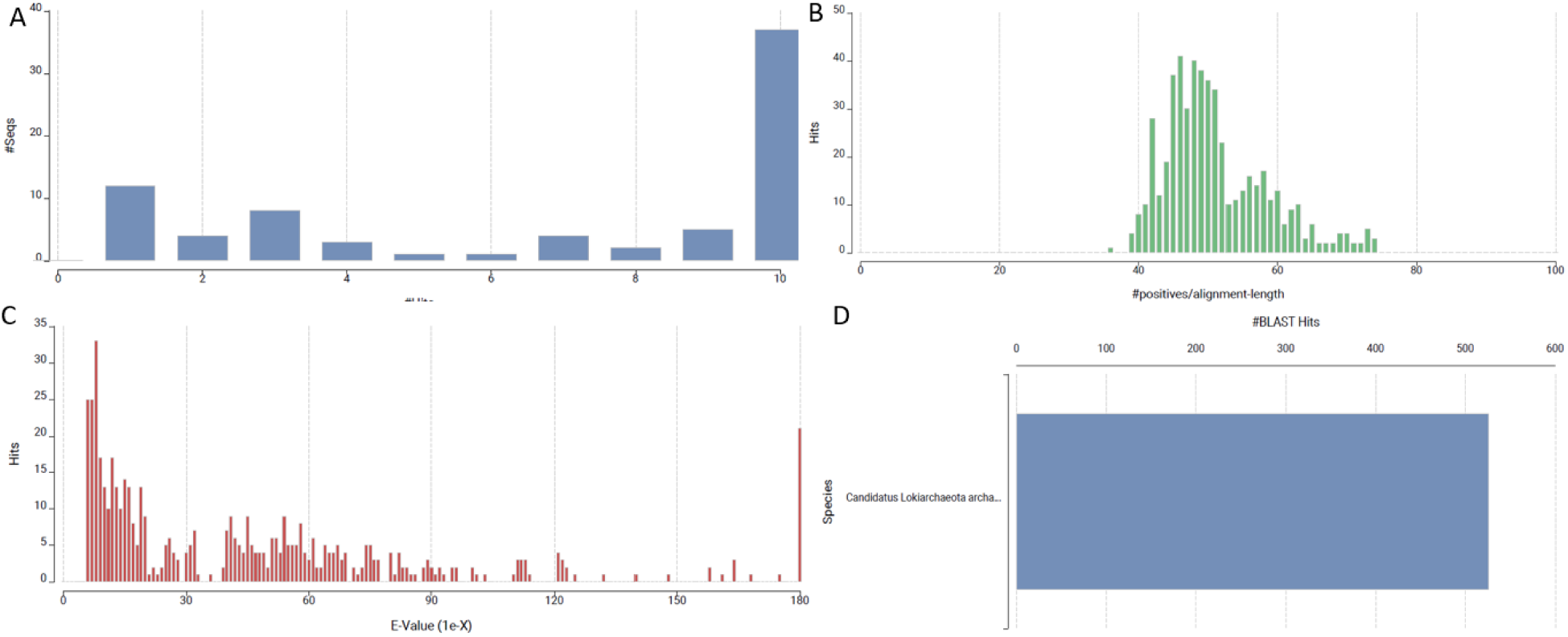
Overview of BLAST analysis results of sleep genes against *Lokiarchaeota* genome. (A) Distribution of hit scores across different ranges, showing higher frequencies at the tail ends. (B) Histogram of the percentage of positive alignments relative to alignment length, peaking around 60%. (C) Distribution of E-values, indicating most hits fall under significant thresholds, with a few outliers at higher E-values. (D) Species distribution of BLAST hits, with Lolrica species accounting for the majority of the matches.

Thus, the fact that the processes connected with sleep have been retained during millions of years of development of at least two highly distinct biological kingdoms - humans and Lokiarchaeota, an archaeal species, emphasizes the importance of the discovery. This conservation implies that core processes of sleep management could be evolutionarily very old and are conserved in many different phyla. Studying proteins such as "GTP-binding protein," "adenosine deaminase," and "serine hydroxymethyltransferase" provides insights into several metabolic pathways that may be linked to sleep. Moving forward, to further elucidate the specific actions of these proteins concerning sleep, a comprehensive analysis of existing research work might be needed. This study shows a varied collection of proteins that may be implicated in sleep regulation, encompassing metabolic processes, protein synthesis, signal transduction, and transport processes. The presence of hypothetical proteins as sleep gene orthologs suggests new possibilities for investigating the molecular mechanisms of sleep and new targets for sleep research.

Distribution of sleep gene hits based on domain, family, and superfamilies Figure 4A illustrates the proportions of various InterProScan domains connected to the genes involved in sleep. Domains are on the y-axis while the number of sequences discovered for individual domains is on the x-axis. Out of all the domains, ‘GPCR, rhodopsin-like’ IPR017452 stands out, with roughly 400 sequences (Figure 4A). Other comprehensible domains include the ‘Ion transport domain’ (40128), ‘Myc-type, basic helix-loop-helix’ (71210), and ‘Receptor, ligand binding area’ (55661). These domains show varying levels of sequences, and they may be related to sleep-related behaviors. Moreover, the domain ‘Period circadian protein’ (IPR048814) and ‘Period circadian-like protein’ (IPR022728) are highlighted since the molecule plays an important role in the control of circadian rhythms. It is seen that all the domains identified in the ‘others’ category are infrequent and occur only occasionally. The existence of numerous protein domains demonstrates the number of molecular factors involved in the process of sleep regulation.

**Figure 4.**
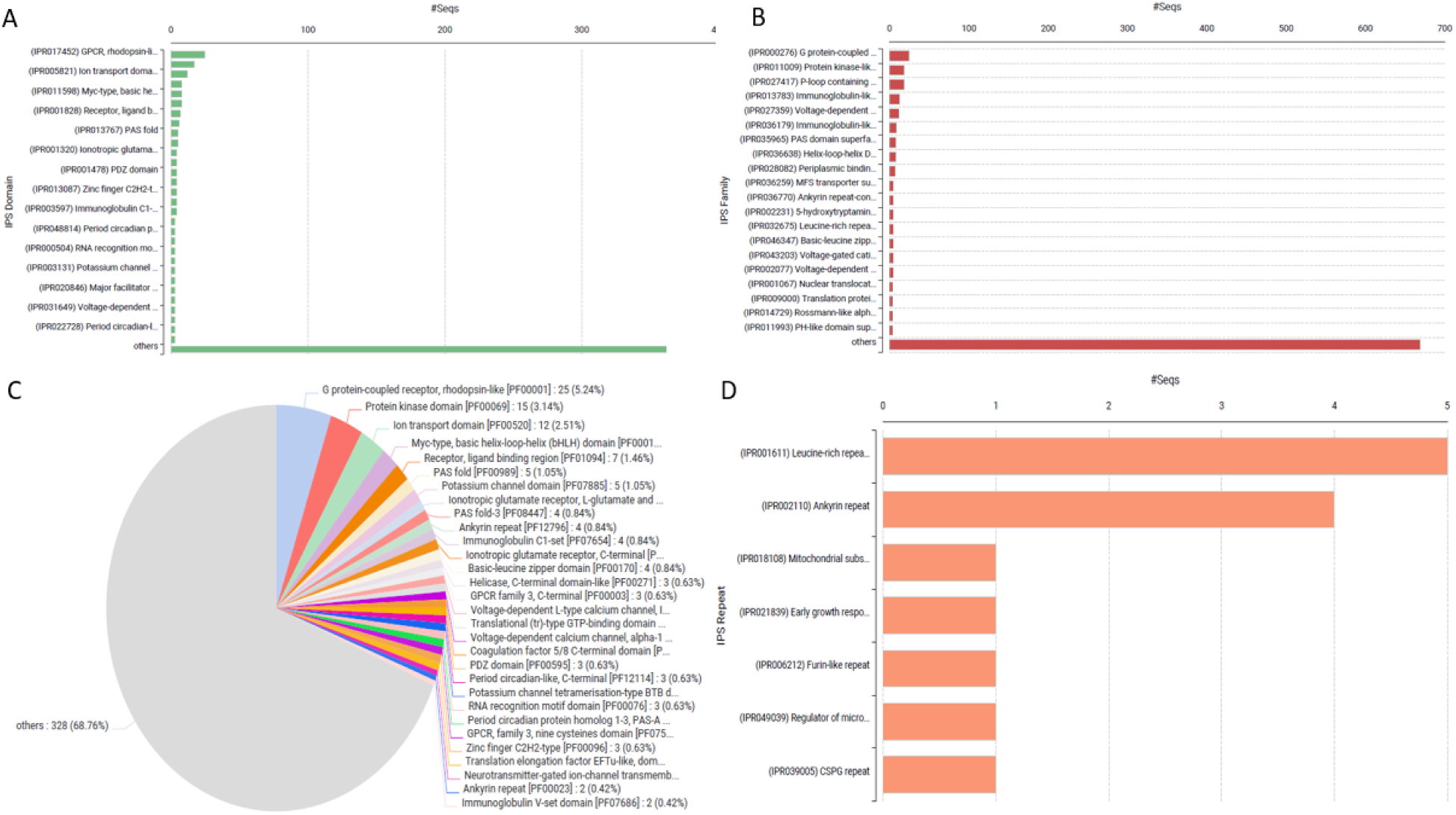
Functional domain and protein family analysis of *Lokiarchaeota* ortholog sequences. (A) Top protein families identified by the number of hits, with GPCR (rhodopsin-like) being the most prominent. (B) Detailed breakdown of specific protein families, showing GPCRs as the dominant family. (C) Pie chart displaying the relative abundance of protein families, with GPCR rhodopsin-like receptors and protein kinase domains representing the largest portions. (D) Top structural or functional motifs based on the number of occurrences, with leucine-rich and ankyrin repeats leading the distribution.

Hence, depending on where their InterProScan IDs are located, all Pfam domains associated with sleep genes are grouped in the following databases (Figure 4B and 4C). The most abundant domain is ‘G protein-coupled receptor, rhodopsin-like’ [PF00001], which accounts for 5% of the structures. 24% of the total. After that, the ‘Protein kinase domain’ with the code PF00069 contributes to a total of 3.9%, the third is the ‘Ion transport domain’ [PF00520] which composes approximately 2.51%. Other considerable domains include ‘Myc-type, basic helix-loop-helix (bHLH) domain, Receptor, ligand binding region’ [PF01094], and ‘PAS fold’ [PF00989], which also contributes to some extent. The ‘Others’ sub-group can be deemed the most dominating formation, as it comprises 68. Logically, the less frequent the domain, the more diversified it is. The part of the total amount of work occupied by these often extremely attractive, yet generally less recurrent domains constitutes 76% belonging to protein families. The detection of varied molecular functional domains in the dataset points towards the fact that sleep regulation entails the many different biological processes that can be deduced from the many Pfam domains.

Figure 4D shows how many repeats of InterProScan are correlated with the genes related to sleep. The ‘Leucine-rich repeat’ (IPR001611) is the most popular repeat followed by the ‘Ankyrin repeat’ (IPR002110). These repetitions are observed in many sequences, and it appears that they are significantly engaged in proteins that are important for sleep regulation. Other repeats include Mitochondrial substrate carrier family repeat (IPR018108); Early growth response protein repeat (IPR021839) and Furin-like repeat (IPR006212) are less common and are found in a relatively smaller number of sequences. It also strengthens the variety of structural motifs, which could participate in molecular underpinnings of sleep regulation, though in lesser amounts are present in the form of several repeating patterns.

### Gene ontology and Enzyme code distribution

The provided dataset shows the 20 most associated GO terms to the genes concerning sleep identified through GO level 20. Figure 5, named "GO Distribution by Level (2) - Top 20," classifies the GO terms into three categories: Actually, it is divided into three categories namely: Biological Process (BP), Molecular Function (MF), and Cellular Component (CC). It demonstrates how often sequences associated with each term appear; phrases like “cellular process,” “biological regulation,” and “regulation of biological processes” are underlined as relevant to the BP category as well as “metabolic process.” In the case of the MF group, the most frequently used terms are “catalytic activity” and “binding” while in the CC category, “cellular anatomical entity” can be observed.

**Figure 5.**
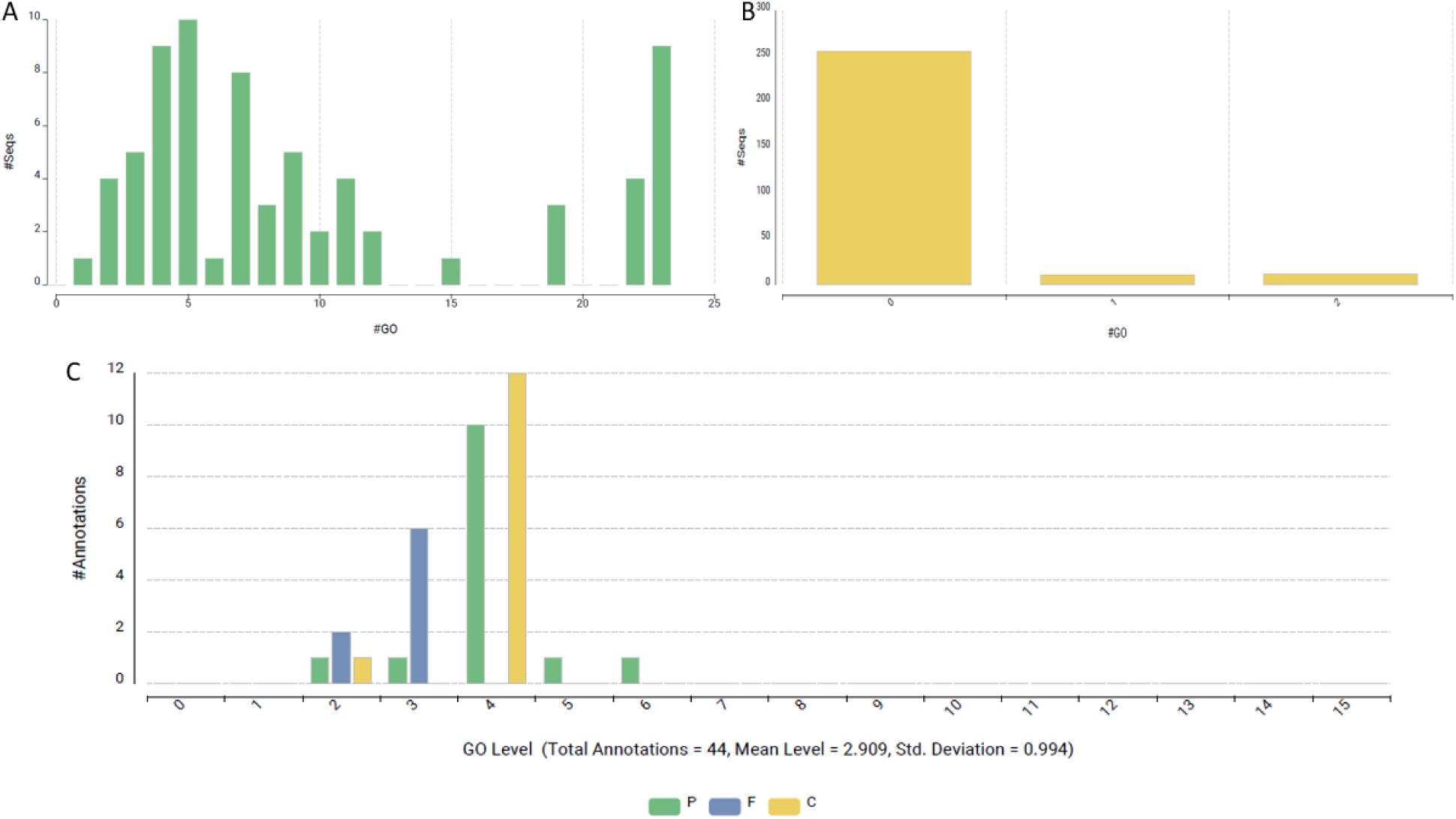
Gene Ontology (GO) term distribution across biological processes, molecular functions, and cellular components of *Lokiarchaeota* ortholog sequences. (A) Frequency of annotations per GO term showing diverse distribution patterns. (B) Dominance of annotations at GO level 0, with significantly fewer terms at higher levels. (C) Breakdown of annotations across GO levels, with categories represented by biological processes (P), molecular functions (F), and cellular components (C). Total annotations amount to 44, with a mean level of 2.909 and a standard deviation of 0.994.

Figure 6A with the label “GO Distribution by Level (3) - Top 20” shows a more detailed division of the GO terms. It gives more focus on specific activities and actions included in the BP such as “regulation of cellular process,” “organic substance metabolic process,” and “primary metabolic process.” Within molecular functions, the term “transferase activity” is important. The cellular component (CC) category entails stress on the terms such as ‘cytoplasm,’ and ‘intracellular anatomical structure.’ Although the numbers of nodes and edges are comparable, the kind and complexity of GO keywords associated with sleep-related genes are illustrated in Figure 6B.

**Figure 6:**
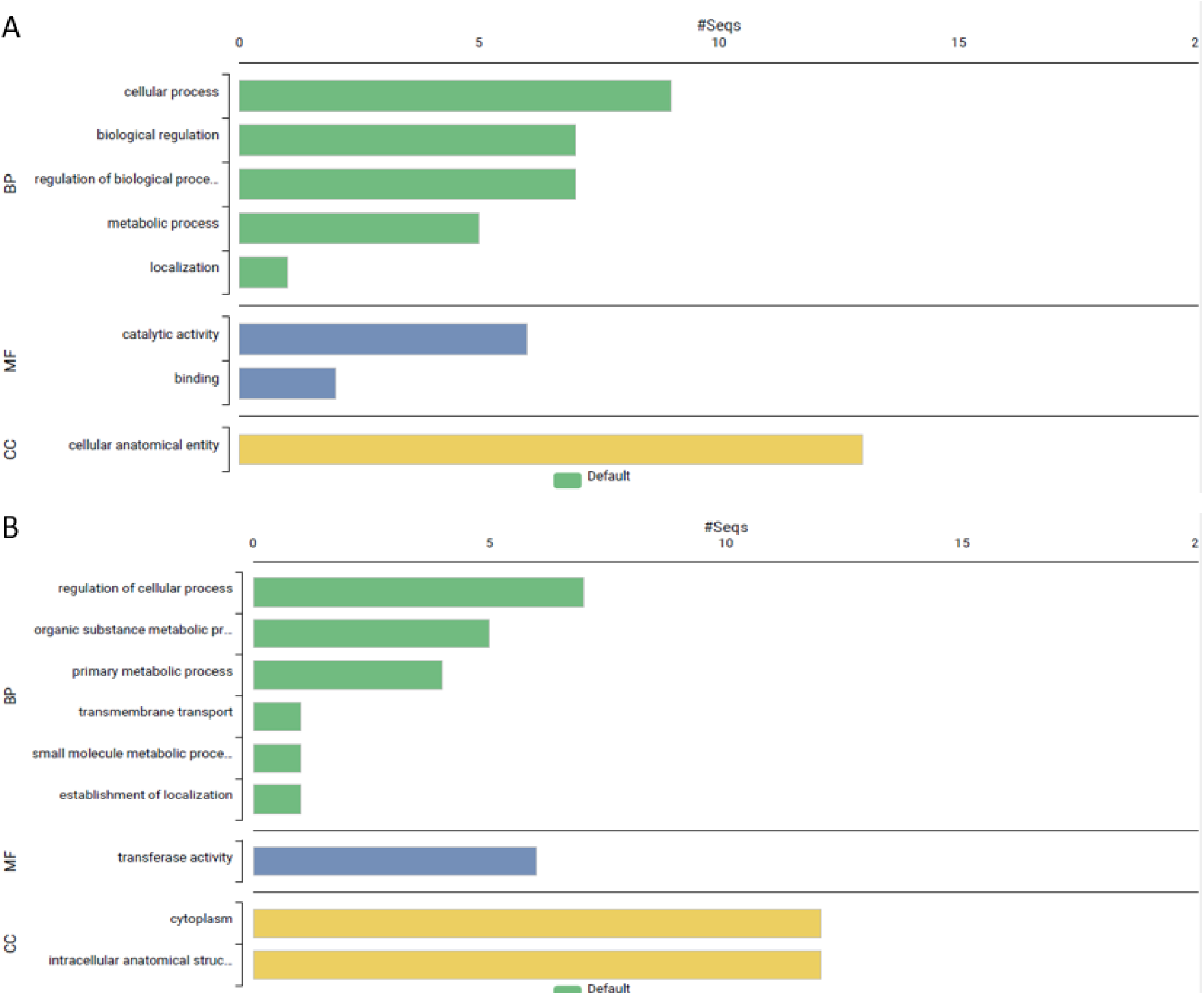
Gene ontology (GO) enrichment analysis results of *Lokiarchaeota* ortholog sequences. (A) and (B) display the enriched GO terms across three main GO categories: biological process (BP, green), molecular function (MF, blue), and cellular component (CC, yellow). The bar length indicates the number of sequences associated with each term, showing the most enriched terms in each category.

As shown in Figure 7A, the distribution of enzyme codes (EC) regarding the transferases class of enzymes includes sleep genes in particular. The x-axis presents the various categories of transferases that are subcategorized into acyltransferases, enzymes that transfer nitrogenous groups, and enzymes that transfer phosphorus-containing groups. The y-axis shows the occurrence of the sequences related to each subclass. Figure 7B shows that, in terms of sequence, acyltransferases and the enzymes transferring a phosphorus-containing group are more frequent than the enzymes transferring a nitrogenous group. This distribution provides useful data concerning the frequency and range of the functions of these enzyme subclasses for genes containing an essential component of sleep.

**Figure 7.**
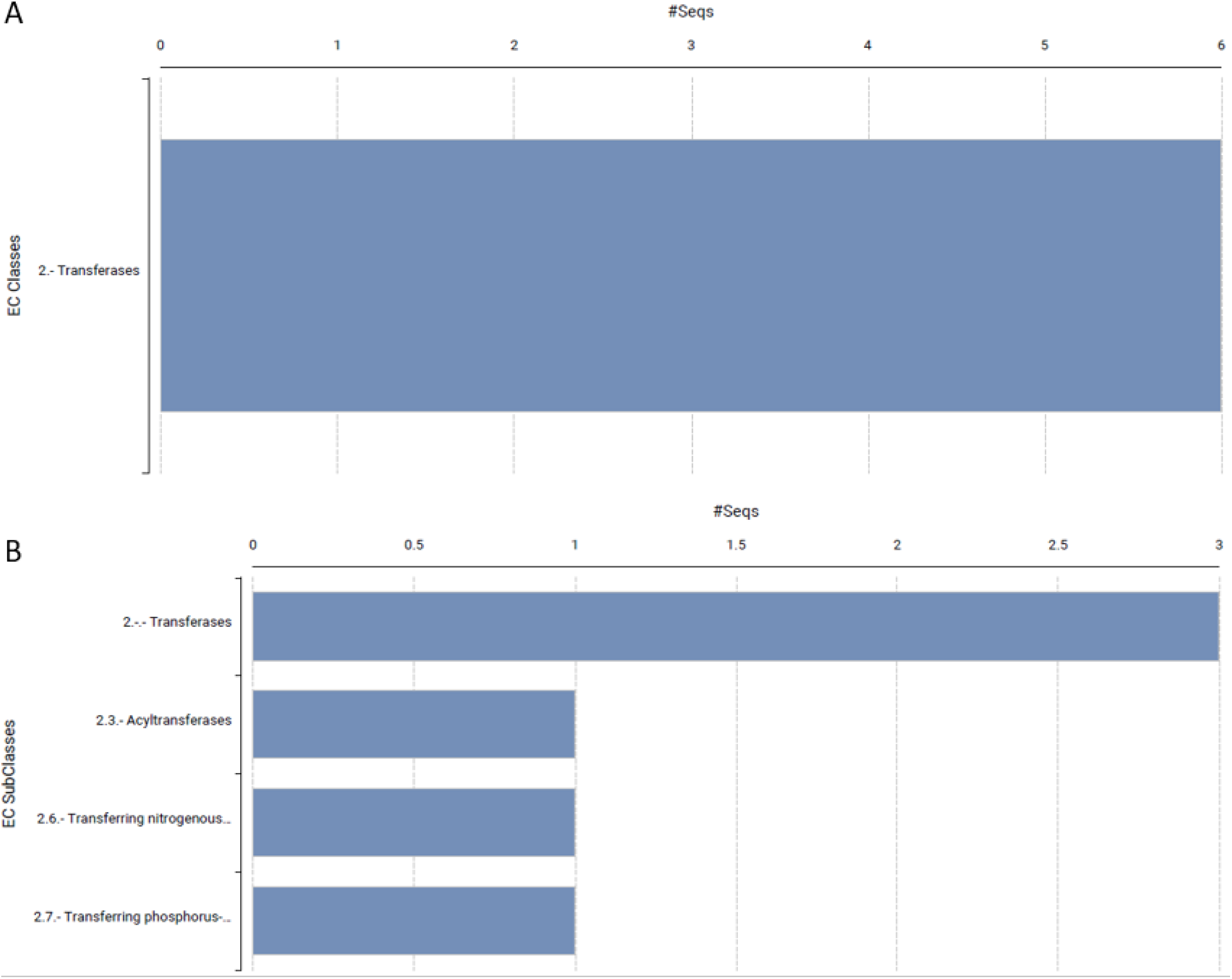
Enzyme classification (EC) analysis results. (A) and (B) show the distribution of enzyme classes and subclasses. (A) displays the number of sequences classified under the "Transferases" category. (B) provides further breakdown into transferase subclasses, including acyltransferases, enzymes transferring nitrogenous groups, and those transferring phosphorus-containing groups. The bar length represents the number of sequences associated with each class or subclass.

### Pathway Analysis

Figure 8A shows the distribution of genes linked to three primary functional categories: Systems: Metabolism, Cellular Processes and Signalling, and Information Storage and Processing. The x-axis shows the percentage of the genes while the y-axis shows the functional categories of the genes. The first and the broadest category is “Cellular Processes and Signalling” which represents the biggest percentage, 34.75%. The term ‘Metabolism’ comes next with approximately 25%, meaning that the pathways of metabolism are an important element within the investigated area. The “Information Storage and Processing” group accounts for approximately 15 percent of the general aggregate of genes and involves the genes necessary for the upkeep and revision of genetic information. It reflects the distribution of cellular activities and signaling pathways as the most significant compound, while metabolism and information processing also play a crucial role in genes’ overall functions.

**Figure 8.**
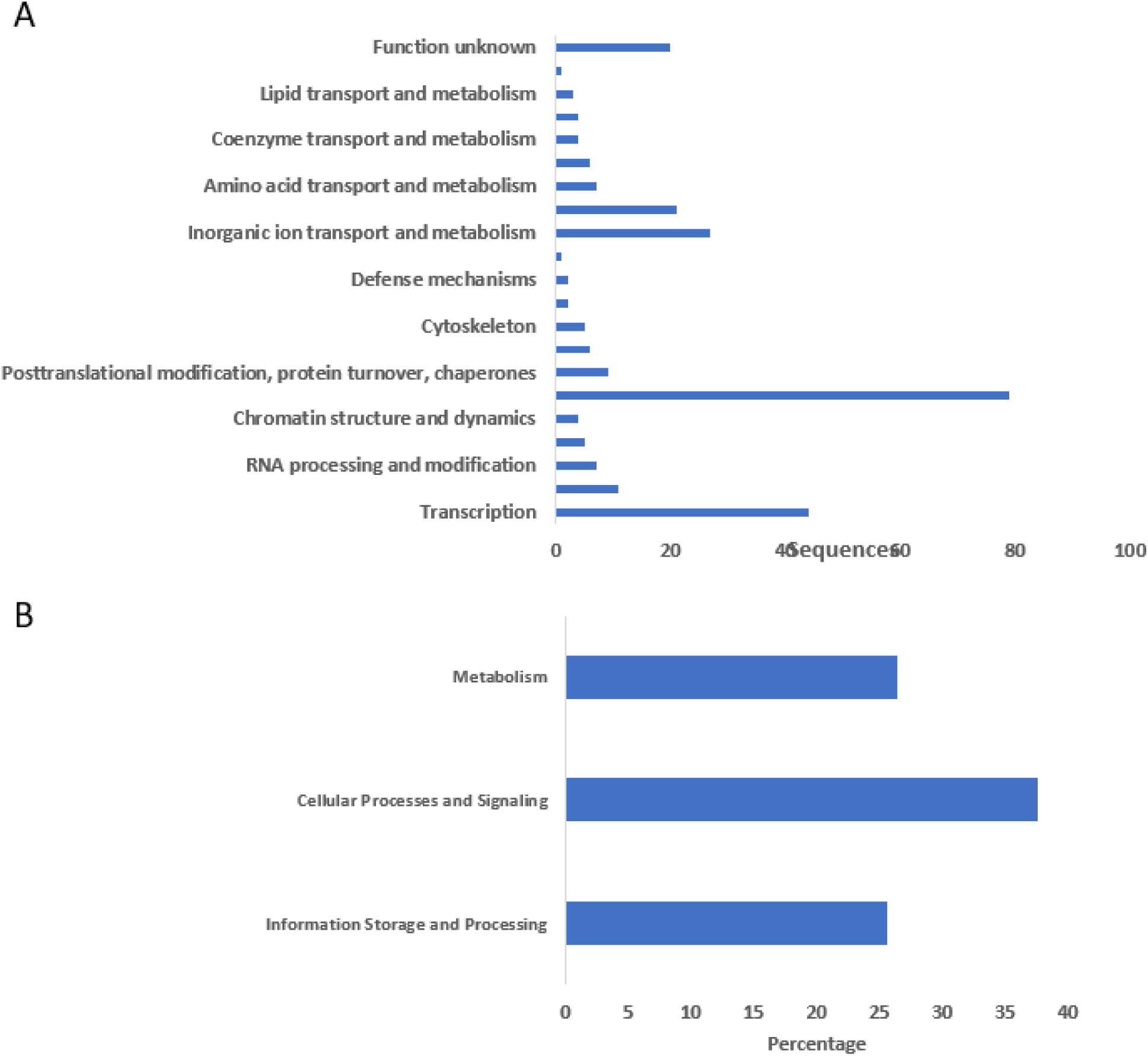
Functional annotation and classification of sequences. (A) Distribution of sequences across various functional categories, highlighting prominent groups such as RNA processing and modification, transcription, posttranslational modification, and chromatin structure dynamics. The "function unknown" category contains sequences that lack known functional annotations. (B) Categorization of sequences into broader functional categories, showing the percentage of sequences involved in metabolism, cellular processes and signaling, and information storage and processing.

Figure 8B shows how various categories of functions are divided in terms of gene sequences. On the X-axis, the numbers represent the number of sequences and on the Y-axis; the letters represent the functional categories. The largest number of sequences is in the category “Signal transduction mechanisms” with more than 80 sequences which can be associated with intensive participation in signal reception and response. The term “transcription” is linked to approximately 40 series, which proves its relevance to the regulation of gene expression. There is also a noticeable representation of categories such as “Inorganic ion transport and metabolism” and “Energy production and conversion.” The “Other” categories, namely, “Function unknown,” “Posttranslational modification, protein turnover, chaperones,” and “Translation, ribosomal structure,” also present moderate levels of sequence counts, hence, diverse but seemingly less functional. Some examples include ’Secondary metabolites biosynthesis, transport and catabolism’, ‘Defence mechanisms’, and ‘Extracellular structures,’ many of which contain few sequences, and could be described as specializing in some function or rare in the genome. The distribution of these cellular processes shows that the processes of signal transduction and transcription are more dominant compared to the other functions.

## Discussion

Most animals sleep, according to extensive exploration of a wide range of phyla with diverse species that differ in shape, size, and other distinctive lifestyle traits. This shows that sleep is a ubiquitous behavior that has been conserved over millions of years of natural selection.

However, its origin, primordial nature, and core function have not been deciphered yet. Some scientists even challenge such a notion and believe sleep may not have any such core function(s) (Geissmann et al, 2024), and consequently has no adaptive value (Gamundí et al, 2005). This may be consistent with Gould and Lewontin’s theory ("Spandrels paper") that, rather than direct adaptations, many higher functions of the human brain are an accidental side effect of natural selection (Gould et al, 1997). Such co-opted traits have sometimes been referred to as exaptation (Gould & Vrba, 1982).

Over the decades, sleep across phylogeny has been investigated (Anafi et al, 2019; Hartse, 2011; Lesku et al, 2009; Capellini et al, 2008; Allada & Siegel, 2008; Lesku et al, 2006; Siegel, 1995; Campbell & Tobler, 1984). Since sleep is believed to be a vulnerable state, it has been shown that, based on behavioral and electrophysiological correlates, some functional characteristics, like adaptation and survival in a particular environmental niche, are found to be retained from an ancestral sleep state (Aime & Adamantidis, 2022; Reinhardt et al, 2019; Siegel, 2012, 2009).

The current findings arising from Lokiarchaeota are rather interesting and puzzling at the same time. What would an earliest known member of the Asgard superphylum exhibit orthologs of sleep genes? Sleep as a behavior may have been shaped over millions of years of slow evolution by several divergent or convergent processes within a particular animal group, which were then impacted by selective forces from the environment and evolutionary bottlenecks. For instance, one of the earliest biologically active signaling molecules in nature is melatonin, also referred to as the "hormone of darkness" and found in a virtually universal range of animals (Pandi-Perumal et al, 2006). It has been suggested that this chemical immediately developed from its antioxidant characteristics. Nighttime especially causes an upregulation of genes related to phototransduction and melatonin production (Tosches et al, 2014).

A study of the molecular and genetic substrate of hormones may have value as a “common language” for analyzing sleep behavior across species that are only distantly related. This may ultimately permit a reconstruction of the “Tree of Life”, first mentioned in the Bible to denote the oneness of all life, and then popularized by Darwin as a metaphor to guide research in evolutionary biology. Until recently, phylogenies were only built using morphological data.

However, discoveries in genetics and molecular biology may enrich our understanding of these phylogenies and reveal further insights into how they are interrelated.

The term phylogeny refers to the study of evolutionary relationships between species using treelike diagrams as representations. There are various limitations to phylogenetic studies (Hao & Xiao, 2020). For instance, it is challenging to determine the evolutionary relationships between different animals because the living world demonstrates incredible diversity in terms of species context. Second, even closely related species do not often appear to be extremely uniform. Third, different biochemical and molecular characteristics may be present in species with comparable phenotypes.

However, the advent of fast, efficient, and robust technological and methodological advances has brought with it several powerful analytical tools. These include processes such as amplification, sequencing, and analytical techniques such as polymerase chain reaction (PCR) (Mullis, 1990a, 1990b; Saiki et al, 1985), quantitative PCR assays (Higuchi et al, 1993), Sanger sequencing (McCombie et al, 2019; Mardis, 2011; Sanger et al, 1977), and molecular phylogenetics. This latter development has given way to phylogenomics, a field in which genome-scale data from multiple samples can be obtained simultaneously at a rapid speed and resolution for a significantly lower cost.

Scientists have studied and discussed the cellular and molecular nature and components of sleep regulation and function over the years. They couldn’t, however, look beyond certain points in their research. As there are obstacles to identifying the primordial nature and function of sleep, we are the first group in the world to ‘sink into the deep ocean approach’ so to speak, and investigate sleep using a combination of phylogenomics and bioinformatics. We believe this approach will be fruitful. Our phylogenomics model offers several advantages. For instance, it reduces noise in data and is highly efficient as it can evaluate a large number of orthologous loci in a short amount of time (Steenwyk & King, 2024).

To summarise, we expected that Loki’s phylogenomic analysis would reveal some genetic relatedness and link between the distantly related taxa, unlike physical traits. Phylogenomics can demonstrate how species are linked to and have developed from one another and the historical characteristics that species have shared. On the other hand, phenotypes in the body may diverge after evolution has passed, but sequences are better conserved.

### Constraints in prior and current approaches in sleep research in elucidating sleep function

There are several limitations in the research tools that are currently used in the field of sleep medicine and sleep research generally. We will first consider generalized limitations in the sleep field per se followed by specific ones pertinent to the current investigation.

First and foremost, the field of sleep science has a major bottleneck, i.e., the definition problem. Sleep is not a unitary state and there is no consensus regarding exactly what it is. Sleep is defined by either electroencephalographic (EEG) criteria (as in the case of humans) or by its behavioral characteristics (in lower animal forms). In humans, sleep has been further divided into two major sleep states namely, non-rapid eye movement (NREM) sleep and rapid eye movement (REM) sleep. The NREM sleep state is further divided into N1, N2, and N3 sleep stages. The functions of these sleep states vary. At a macro level, several variations in sleep behavior have been reported. For example, age and sex differences, variation in sleep duration in animals in the wild vs. captivity, sleeping sites, and so on.

There are other inconsistencies as well. Electroencephalography architecture of Dolphin’s brain, for instance, shows unihemispheric slow wave sleep (USWS) pattern, suggesting that they may be able to swim continuously even while they are sleeping (Lyamin et al, 2008; OLEKSENKO et al, 1992; Mukhametov, 1987). Moreover, brain maturation and development are another suggested role of sleep (Jiang, 2020; Marks et al, 1995; Rao & Chakrabarti, 2004; MIRMIRAN & VAN SOMEREN, 1993). However, it has been noted that during the postpartum period, sleep is greatly reduced in newborns and their mothers of bottlenose dolphins (Tursiops truncates) and killer whales (Orcinus orca) and exhibit virtually no sign of typical sleep behavior (e.g., immobility, unilateral eye closure, surface rest, bottom rest, quiescent hanging, slow stereotypic circular swimming) (Lyamin et al, 2005). This defies the presumed immobility and unresponsiveness that are currently thought to be behavioral characteristics of sleep.

The current ambiguity surrounding the definition of sleep might not have been a major concern at that time, as the research was primarily focused on understanding the nature and physiology of sleep by studying humans and a few key model organisms. Later, interest focused on the ontogeny and phylogeny of sleep. Then, scientists discovered that lower animal forms barely fit the same definitional criteria as humans. Additionally, more recent research has suggested that central brain architecture is not needed, as evident from Cassiopeia and Hydra, which do not possess these structures (Kanaya et al, 2024; Nath et al, 2017).

Furthermore, there are limitations to phylogenetic approaches (Hillis, 1995). At least one of our fundamental presumptions needs to shift for us to find something genuinely novel. That’s where phylogenomics comes into play.

In sleep and circadian clock research, simpler organisms provide excellent genetic models (Pandi-Perumal et al, 2024; Pandi-Perumal, 2010). At a molecular level, unquestionably, genetic approaches have great strength. They cannot, however, fully simulate all facets of mammalian sleep due to the simplicity of the model animals which makes them genetically manipulable (Rattenborg et al, 2008). As a counterargument, it could be objected that many biological networks are somewhat simple. The behavior of biological networks that appear to be inexplicable may eventually be understood thanks to the simplification of concepts. In biology, simplicity has been highlighted to support the idea that broad principles can be elucidated. It is hard to see how one could ever understand biology at the level of a single cell, tissue, or organism without such concepts (Alon, 2007).

Sleep is an enigmatic state, which is observed among phylogenetically distant organisms. However, an organism’s sleeping patterns may have been molded by a combination of basic biological necessities, uncommon and harsh habitats, social habits, geophysical conditions including light/dark (LD) cycle, and adaptive or maladaptive selection pressure. Because of their glaring inconsistencies, current theories and hypotheses are incapable of sufficiently accounting for all these external elements (Pandi-Perumal, 2010).

The literature encompasses various approaches and challenges related to phylogenomics, which involves examining evolutionary relationships through genomic data. Nishihara et al. (2007)(Nishihara et al, 2007) provide a comprehensive review of the strengths and weaknesses of phylogenomic analyses in identifying the eutherian tree, emphasizing data quality and model selection in the reconstruction process. Steenwyk and King (2024) (Steenwyk & King, 2024) emphasize the potential and limitations of synteny in phylogenomic trees, noting its capacity to reveal evolutionary relationships while cautioning against certain weaknesses of the approach. Ultraconserved elements (UCEs) are innovative markers that were introduced by McCormack et al. (2012) (McCormack et al, 2010), which have proven value when integrated with species tree analysis. This approach has enhanced the understanding of the phylogeny of placental mammals and has demonstrated the effectiveness of UCEs in addressing evolutionary challenges. In summary, these studies highlight both the advancements and constraints of genomic analysis for deducing the nature of phylogenies.

So far, the quantitative polymerase chain reaction (QPCR) analysis has been performed to monitor the growth of the organism culture using the amplification of 16S rRNA genes (Imachi et al, 2020). However, the validation of the expression of sleep-related genes associated with the organisms has not been explored so far due to the complexity associated with the species. Our data will be the stepping stone for validating the sleep-related genes in this species.

Despite several of these limitations, this is the first study that identifies the presence of human sleep genes in Loki. What does it signify? It has important ramifications for our knowledge of the evolution of sleep in general, and possibly, its genetic roots. From the evolutionary perspective, despite vast evolutionary time scale differences, these genes are conserved. This implies that evolution has maintained its significance and functional roles. For example, these genes associated with certain diseases often participate in certain basic biological processes.

Hence, exploring these genes in simpler organisms helps to decipher the origin of sleep and sleep disorders. Therefore, smaller organisms serve as valuable tools for studying the functions of sleep or sleep disorders in a simpler biological context. Such research can elucidate the basic mechanisms by which these genes operate, interact, regulate, or dysregulate certain biological functions, which eventually lead to disease in humans.

Research on sleep and sleep disorders genes in ancient organisms can provide valuable information for drug development and scientific research. Knowing the conservation and evolutionary background of genes linked to disease could lead to the discovery of novel therapeutic targets. Researching disease genes in simpler organisms is a valuable tool for comparative genomics and phylogenomics, which compares genetic sequences and structures between various species, and across phyla. This method aids in locating conserved sequences that are essential to gene function and susceptibility to illness. In conclusion, the importance of human illness genes in other simpler organisms can be attributed to their understanding of the origin of sleep, sleep disorders, evolutionary conservation, fundamental biological processes, and possible use in the development of novel therapeutics and treatment modalities.

## Conclusion

"Nothing in biology makes sense except in the light of evolution"

- Dobzhansky, 1973

In his famous dictum, Dobzhansky emphasized the need for an evolutionary approach in biological sciences. In the current work, we have used both phylogenomics and bioinformatics to an unresolved biological problem. The scientific investigation of sleep is a very dynamic and fascinating area of study due to the rapid technological advancements in genetics. There are considerable challenges, concerns, and opportunities ahead in every level of investigation.

Based on the phylogenomics and bioinformatics approach, this research illustrates the scope of present study findings and offers a glimpse of what sleep science may hold in store for the future.

Sleep is a pervasive, conspicuous, ubiquitous, and adaptive behavior in the animal kingdom yet there is no universal definition of sleep based on the characteristics that have been investigated to date. Despite decades of investigation, the exact nature, purpose, and function are unknown. As to whether sleep serves a single essential purpose (i.e., core or universal vital or primordial function), the answer continues to be debated. Why sleep is necessary and maintained by evolution is yet unknown. From an evolutionary standpoint, distant and unrelated organisms sleep. Many scientists believe that phylogenetic approaches would help to unravel some of these mysteries. Although such work has advanced the sleep field to some extent, there are certain bottlenecks (discussed earlier) as it has limitations of its own. Therefore, it is not a complete solution for the elucidation of sleep function.

The phylogenomics approach to sleep science is rather new, however, it is paramount to comprehend the function of sleep. Although the basic function of sleep remains mostly unknown, combining phylogenomics and bioinformatics research on a range of species may help to narrow the field of inquiry. In this regard, our study of sleep in a most disparate species - like Loki - has unveiled some surprising results. This study’s principal findings include identifying a conserved molecular pathway in the distant species, Loki, which appears to regulate essential sleep characteristics. This pathway, encompassing genes associated with energy metabolism and cellular repair, suggests that sleep may be a crucial physiological mechanism for cellular maintenance. These findings corroborate the concept that sleep is an evolutionary conserved biological function while also elucidating new features of the methods by which evolutionarily distinct creatures have developed this process. The findings align with previous assumptions concerning the restorative and protective functions of sleep and comprehensive cellular maintenance, including gene protection. In the end, we hope that by disseminating our findings, other scientists will be inspired to look for clues elsewhere, especially when they have unresolved scientific problems. In the field of sleep, there are ongoing controversies, inconsistencies, and unknowns that are yet to be worked out.

Given the critical benefits of sleep in humans, It’s imperative to fill in some of the knowledge gaps. It is hoped that our work will help advance our understanding of why humans sleep.

## Contact details of all authors

Seithikurippu R Pandi-Perumal –

Konda Mani Saravanan -

Sayan Paul -

David Warren Spence –

Saravana Babu Chidambaram -

## Acknowledgments

We are grateful to Dr. Fredrik Sahlström for the supply of the location map of Loki’s Castle (figure 1). Due to the article’s limited emphasis and other circumstances, such as page constraints, it was not possible to reference all relevant articles on this topic. As academics, we frequently must make such difficult decisions when multiple remarkable studies are excluded for editorial reasons.

## CRediT authorship contribution statement

SRP; SBC: Conceptualization and Methodology; SRP; KMS; SP: Data collection and review, KMS; SP: Software, KMS, SP: Visualization; SRP; KMS; SP: Preparation of original draft; SBC: Input to the original draft and critical review, interpretation, and editing of the final version of the manuscript; SBC: supervision. Prior to submission, all authors (SRP; KMS; SP; SBC) have contributed to review, editing, read, and agreed to the submitted version of the manuscript.

## Author Agreement Statement

All authors confirm that the paper is their original work that has not previously been published nor is presently being considered for publication anywhere. Each author expressly declares that they contributed equally and intellectually, read, evaluated, and accepted the final version of the paper, and agreed to be included as co-authors per ICMJE guidelines. Each of the authors has approved the author sequence and the corresponding authors.

## Ethical Statement

This study does not contain any work involving animals or human participants performed by any of the authors. Hence, no IRB approval was necessary for this work.

## Data availability

The study authors created a dataset, which is being utilized for the ongoing prospective phylogenomics investigation. Hence, except for the data shared in this work, complete source data sharing may not be possible.

## Code availability

Not applicable

